# Accessory genome dynamics and structural variation of *Shigella* from persistent infections

**DOI:** 10.1101/2020.09.28.316513

**Authors:** Rebecca J. Bengtsson, Timothy J. Dallman, Claire Jenkins, Hester Allen, P. Malaka De Silva, George Stenhouse, Caisey V. Pulford, Rebecca J. Bennett, Kate S. Baker

## Abstract

Shigellosis is a diarrhoeal disease caused mainly by *Shigella flexneri* and *Shigella sonnei*. Infection from *Shigella* is thought to be largely self-limiting, with short- to medium- term and serotype-specific immunity provided following clearance. However, cases of men who have sex with men (MSM) associated shigellosis have been reported where *Shigella* of the same serotype were serially sampled from individuals between 1 to 1862 days apart, possibly due to persistent carriage or reinfection with the same serotype. Here, we investigate the accessory genome dynamics of MSM associated *S. flexneri* and *S. sonnei* isolates serially sampled from individual patients at various days apart. We find that pairs likely associated with persistent carriage infection and with smaller single nucleotide polymorphism (SNP) distance, demonstrated significantly less variation in accessory genome content than pairs likely associated with reinfection and with greater SNP-distance. We also observed evidence of antimicrobial resistance (AMR) acquisition during persistent *Shigella* infection, specifically the gain of extended spectrum beta-lactamase genes in two pairs associated with persistent carriage. Finally, we explored chromosomal structural variations and rearrangements in seven (5 chronic and 2 reinfection associated) pairs of *S. flexneri* 3a isolates from a MSM-associated epidemic sublineage, which revealed variations at several common regions across pairs. These variations were mediated by insertion sequence (IS) elements which facilitated plasticity of genetic material with a distinct predicted functional profile. This study provides insight on the variation of accessory genome dynamics and large structural genomic changes in *Shigella* during persistent infection.

**Importance:** *Shigella* spp are Gram-negative bacteria that are the etiological agent of shigellosis, the second most common cause of diarrhoeal illness globally, particularly among children under the age of 5 in low-income countries. In high-income countries, an alternative transmission pathway of sexually transmissible disease among men who have sex with men (MSM) is emerging as the dominant presentation of the disease. Within MSM we have captured prolonged infection and/or recurrent infection with shigellae of the same serotype, challenging the belief that Shigella infection is short-lived, and confers homologous serotypic immunity. Using this recently-emerged transmission scenario we comprehensively characterise the genomic changes that occur over the course of individual infection with Shigella and uncover a distinct functional profile of variable genome regions in these globally important pathogens.

## Introduction

Shigellosis is a faecal-orally transmitted disease that is characterised by dysentery and severe colitis. The causative agent is the Gram-negative bacteria *Shigella* spp. *S. flexneri* and *S. sonnei* contribute to the greatest disease burden of shigellosis globally, are among the leading cause of moderate-to-severe diarrhoea in children under the age of five in low-income countries (1). In high-income countries, cases are often linked to foreign travel and can be sexually transmitted among GBMSM, evidenced by an increase in the number of domestically-acquired infection cases among adult males (2–4).

The current recommended treatment for shigellosis is ciprofloxacin. However, *Shigella* spp with chromosomal mutations in the Quinolone Resistance Determining Region (QRDR) conferring resistance to fluoroquinolones, are now widely geographically distributed (5, 6) and have been reported in MSM-associated outbreaks (4, 7, 8). Genomic epidemiological analysis has previously shown that horizontal acquisition of a single azithromycin resistance plasmid, pKSR100, facilitated the epidemic emergence of MSM-associated shigellae in 2012 and enhanced its spread (9). Over the recent years, increase in MSM shigellosis in the UK has been attributable to a novel *S. sonnei* clade exhibiting ciprofloxacin and macrolide resistance, conferred by triple QRDR mutations and the acquisition of pKSR100, respectively (10). Thus, MSM-associated shigellosis is an emerging problem that is intimately associated with increasing AMR (2).

In addition to AMR, coinfection with HIV may lead to further complications and can be a risk factor for sustaining ongoing transmission within the MSM community (11, 12). *Shigella* infection is typically self-limiting (infection time ranging between 1 to 4 weeks) and following clearance immunity is acquired against subsequent infection with the homologous serotype (13). The conferred length of protection is thought to last approximately 5 months to 2 years (14, 15). However, persistent infection has been reported among MSM and coinfection with HIV could be a contributing factor, altering individual immune statuses and causing prolonged infection times, relapse or re-infection of immunocompromised individual with the same serotype (2, 16). Due to its rarity, little is known regarding persistent *Shigella* infection and it remains poorly characterized. The intensification of MSM-associated shigellosis in England over recent years has provided a diverse dataset of *Shigella* isolate pairs serially sampled from individual male patients reporting domestically acquired infection across several serotypes (*S. flexneri* 3a, *S. flexneri* 2a and S. *sonnei*) (17). Previous analysis of these pairs revealed SNP distances between such paired isolates increased with time between sampling and demonstrated patterns of long-term carriage or recurrent infection, with either the same or different serotypes (17).

Here, we extend previous SNP based comparisons among serially isolated *Shigella* pairs (n=58 pairs) and perform detailed comparative analyses to investigate genomic changes in shigellae over the course of infection. We characterise accessory genome dynamics, including the gain and loss of AMR determinants, compare and contrast these changes between pairs that represent long-term carriage to those that arose from reinfection. We then further deepen the study to compare large-scale structural variation across the *Shigella* chromosome through long-read sequencing of a subset of pairs (*n*=7 pairs). In doing so, we generated a high-quality reference genome and publicly accessioned an isolate of a globally important pathogenic *S. flexneri 3a*. We also identified unique functional signatures in variable regions of the chromosome, providing a snapshot into the genome changes that occur over the course of infection.

## Materials and Methods

### Isolates with routinely generated Illumina sequencing data

Whole genome sequencing data of *Shigella* isolates (*n*=116) were generated as part of routine national surveillance by Public Health England (18, 19). Each patient was sampled at two time points ranging from 1 to 1,862 days apart for *S. flexneri* and 1 to 1,353 days apart for *S. sonnei*. In total, the dataset consists of 35 pairs of *S. flexneri* (19 paired *S. flexneri* 2a and 15 paired *S. flexneri* 3a) and 23 pairs of *S. sonnei* (Table 1). All 68 *S. flexneri* and over half of *S. sonnei* (28/46) isolates belonged to previously described epidemic MSM-associated lineages (2, 9). As established by Hester *et al*, we define a pair of isolates serially sampled from the same patient at time points ranging 1 to 176 days with genetic distance ranging between 0 to 7 SNPs, as likely associated with long-term carriage, and pairs of isolates serially sampled between 34 to 2,636 days with genetic distances of 10 to 1,462 SNPs, as likely associated with re-infection (17). Here, we simplify these nomenclatures as ‘carriage associated’ and ‘re-infection associated’ isolate pairs. Using these definitions, the dataset used for analysis comprised of 22 carriage associated and 12 reinfection associated pairs for *S. flexneri*, and 15 carriage associated and 8 reinfection associated pairs of *S. sonnei* (Table 1). Further details regarding individual isolates used in this study and the Sequence Read Archive (SRA) accession numbers are listed in Table S1.

**Table 1.** Number of isolate pairs analysed in the current study, broken down by *Shigella* serotypes and classification as carriage associated or reinfection associated.

| Serotype | Carriage associated pairs | Reinfection associated pairs | Total pairs |
| --- | --- | --- | --- |
| *S. flexneri* 2a | 14 | 5 | 19 |
| *S. flexneri* 3a | 8 | 7 | 15 |
| *S. sonnei* | 15 | 8 | 23 |

### Extension study of S. flexneri 3a isolates including long-read sequenced isolates

Sixteen epidemic sublineage MSM-associated *S. flexneri* 3a isolates (2) were used to determine large structural variation and genome rearrangement of *Shigella* over time. These isolates were serially isolated from eight individuals between 9 to 911 days apart and were sequenced with both Illumina and PacBio technologies (Figure 3A and 3B). For one patient (sampled with 154 days interval), PacBio sequencing was only successful for the earlier isolate. Illumina sequencing for the isolates were generated at the Wellcome Trust Sanger Institute, as previously described (2). For this study, the 16 isolates were revived from the Gastrointestinal Bacterial Reference Unit reference laboratory archives and DNA extracted for long read sequencing as previously described (9). DNA from each sample was sequenced on a Pacific Biosciences RS Sequel at the Centre for Genomics Research at the Institute for Integrative Biology, University of Liverpool.

To facilitate understanding, the isolates and near-complete genome assemblies in this extension study have been abbreviated to meaningful titles to reflect the epidemiology and sequencing technology employed. These names comprise: the number of intervening days between serial isolations, time point (A or B, being earlier and later time points respectively) and sequencing technology (Illumina [I] or PacBio[P]) (Figure 3B). For example, 20BP is the PacBio-sequenced genome of the second isolate taken from a patient whose isolates were sampled 20 days apart. The full key and genome accession numbers are provided in Supplementary Table 2.

### Sequence processing and assembly

Illumina sequencing data was adapter- and quality-trimmed using Trimmomatic v0.38 (20) and draft genomes were assembled using Unicycler v0.4.7 (21). PacBio data was assembled using canu version 1.6 (22) and iteratively polished using SMRT tool (Arrow) version v6.0.0 (https://github.com/PacificBiosciences/GenomicConsensus). This generated genomes with a variable number of contigs (between 3 and 17, mode 6). These draft genomes were re-ordered against the completed reference genome (below) manually using a combination of pairwise all-by-all Basic Local Alignment Search Tool (BLAST) and bedtools v2.27.1 (23).

### Generation of a public isolate and complete genome of an internationally important pathogen

For one PacBio sequenced genome (20BP), three contiguous sequences were generated that corresponded to the bacterial chromosome, virulence plasmid and pKSR100 resistance plasmid. To complete this genome, circularisation at *dnaA* was achieved manually by self-BLAST and removal of inverted repeat regions using bedtools. As this belonged to an internationally important pathogen, the cognate isolate has also been deposited at the National Collection for Type Cultures (NCTC) under accession number xxxxx <awaiting accession>. Complete genome of 20BP was cut and linearized at *dnaA,* this was then used as a reference for downstream analyses.

### Pangenome and pairwise homologous sequence search

All assembled draft genomes were annotated using Prokka v1.13.3 (24) and pangenome analyses were performed using Roary v3.12.0 (25), run without splitting paralogs. To determine gain and loss of genes, pairwise homologous sequence search was carried out using Roary between pairs serially isolated from individual patients at two time points. Accessory genes present in the first isolate and absent in the second were classified as lost, while genes absent in the first isolate and present in the second were classified as gained. To account for variations of gained/lost genes contributed by misassembly and inaccurate annotation, seven synthetic read sets of lengths 36 – 90bp and variable insert sizes (Supplementary Table 3) were generated from each of the complete genomes of *S. flexneri* 20BP and *S. sonnei* Ss046 (GenBank assembly accession: GCA_000092525.1) using the randomreads.sh script from the BBMap package (26). These synthetic read sets were then assembled, annotated and underwent pairwise comparisons (as above). Comprehensive pairwise comparisons were ran among the seven synthetic draft genomes generated from each reference genome. By which, each genome assembled from a particular read length were individually compared to the six genomes assembled at various lengths, generating a total of 42 pairwise comparison for each species.

### Detection of previously characterised accessory genome elements

The presence of genetic determinants conferring AMR were detected using AMRFinder v3.1.1b (27). Plasmids were identified in genome assemblies through screening for plasmid amplicons using PlasmidFinder with >98% sequence identity and 100% query coverage (28). Presence and absence of the pKSR100, pCERC1, spA plasmid were confirmed using short read mapping with BWA mem against the pKSR100 from *S. flexneri* 20BP, pCERC1 from *E. coli* S1.2.T2R (Genebank accession JN012467) and spA from *S. sonnei* Ss046 (Genebank accession CP000641) (29). Mapping of more than >90% sequence coverage across the reference were defined as present. Further mobile elements were identified by BLAST of contiguous sequences using MegaBlast against the NCBI non-redundant database. Phage elements in the 20BP reference genome were predicted using PHASTER (30).

### Core SNP distances and phylogenetic inference

In order to measure the genetic distances between each pair of isolates sequenced from MSM-associated *S. flexneri* 3a epidemic sublineage, pairwise SNP distances were ascertained as previously described (2) with the following exceptions. The reference genome used was 20BP along with its associated virulence and resistance plasmid. The short-read Illumina data was mapped directly and the PacBio draft assemblies were shredded to simulated data of 100bp in length with a 250bp insert size every three bases along a circular chromosome, as previously described (9), before mapping as for Illumina data.

### Structural rearrangements and functional annotation

In order to detect structural variations and genome rearrangement among pairs, the Synteny and Rearrangement Identifier (SyRI) package was used. First, the 14 PacBio assembled draft genomes were reordered against the complete reference genome of 20BP using chroder, part of the SyRI software package (31). Then, using the NUCmer utility, reordered genomes were individually aligned against 20BP reference genome, alignment coordinates generated were then used as input for SyRI to detect structural variation between isolate pairs. The output of SyRI was compared between the two isolates in each pair, common variations (detected in both isolates) suggested inter-isolate variation of the pair with the reference genome, whereas unique variations (detected in one of the isolate pairs) suggested intra-isolate variations. Insertions and inversions detected by SyRI were evaluated by visualizing pairwise comparison of PacBio draft assemblies using Artemis Comparison Tools (32). Mapping of short- and long-reads at regions of intra-isolate variation was performed to confirm duplications and deletions detected by SyRI and verified manually using Artemis visualisation of coverage at the region (33). Where consistency between short and long-read mapping was found, a true biological structural variation between isolate pairs was indicated. However, discrepancies between short and long read mappings may suggest variation introduced by different sequencing technologies or through the difference between revived DNA preparations of the same isolate (Figure 3A). Coordinates of the structural variants identified among the seven *S. flexneri* 3a pairs (according to the location of the 20BP reference genome) were parsed to Circos for visualization (34).

To explore the functional features of the structural variable genomic regions, locations of the variable regions borders were identified along the 20BP chromosome, and genome sequences were manually checked for IS elements, as identified using ISEScan (35). Functional assignment of the Gene Ontology (GO) category for genes in the 20 BP reference chromosome was predicted using RAST (36), which annotates CDS by comparison to the curated FIGfams protein families database (37) and assigns genes into different functional categories.

### Statistical analyses

All statistical analyses were performed using R language v3.6.1. Statistical differences between accessory gene content variation among isolate pair classification groups (i.e. carriage vs. reinfection and data vs. control) were tested using the Mann-Whitney U test (38) using the wilcox.text() function. Linear regression analysis of SNP distance against gene content variation among isolate pairs was performed using the lm() function. The correlation between gene content variation and SNP distance was tested using the Spearman’s rank correlation coefficient using the cor.test() function. Statistical difference in the proportion of genes in each GO category was tested using Chi-square tests with the chisq.test() function and using the raw values.

### Data availability

All data have been deposited in the European Nucleotide Archive under the study accession number PRJEB39785 with individual isolate accessions listed in Table S1.

## Results

### Change in accessory genome over time among carriage and reinfection isolate pairs

Here we defined carriage and reinfection associated pairs based on SNP distance and isolate pair serial sampling time interval, according to previous definitions (see methods). In order to extend our understanding of the accessory genome dynamics during the course of *Shigella* infection, we examined the difference in gene contents between pairs of carriage and reinfection associated isolates. First, we assessed the correlation between SNP distances and gene content variation, which was positive and statistically significant for both species, although the association is stronger for *S. sonnei* (*r* = 0.80, Spearman’s rank correlation coefficient) than *S. flexneri* (*r* = 0.56). (Figure 1). This indicated that gene content variation increases as the genetic distance between a pair of serially sampled isolates increases.

**Figure 1.**
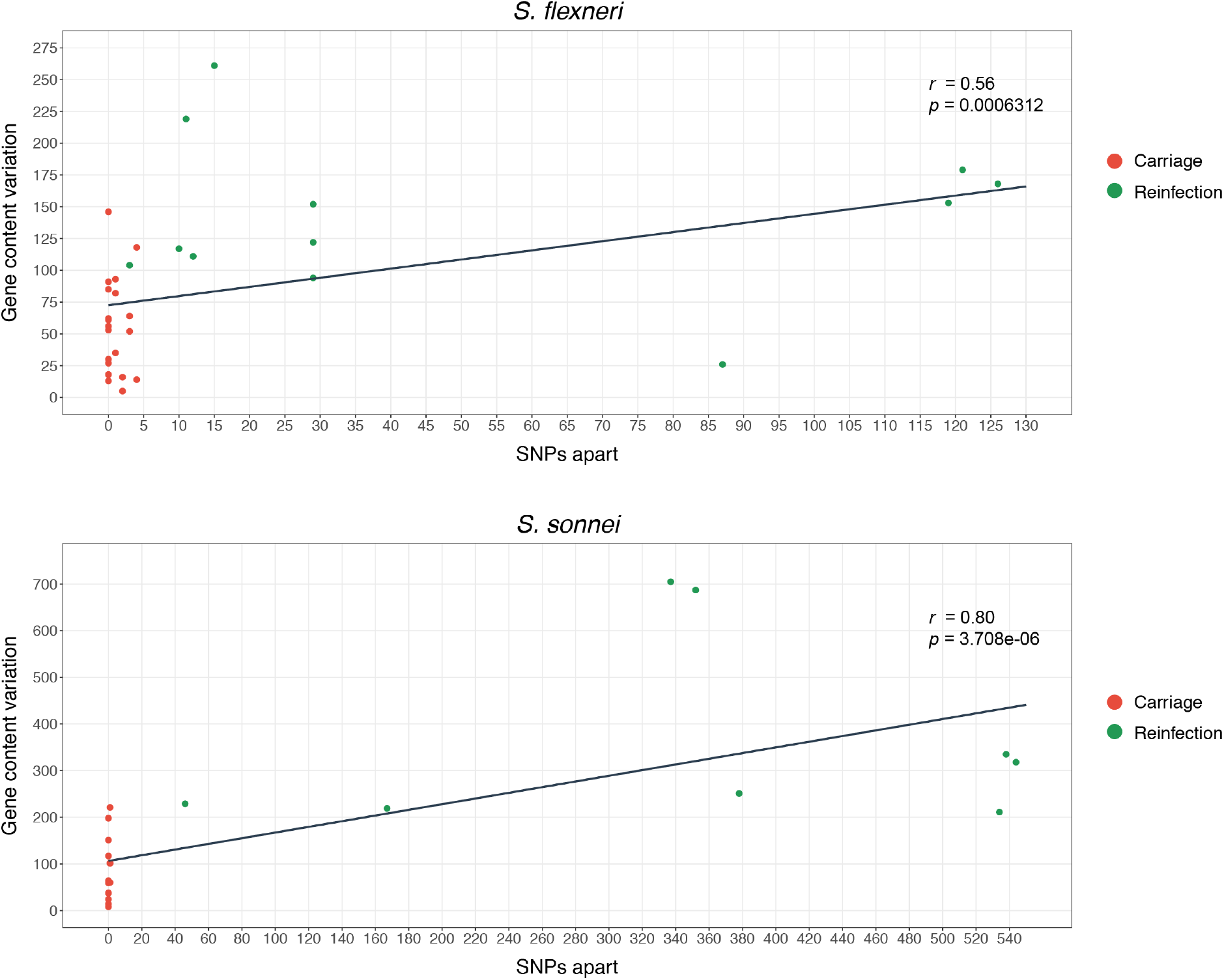
Association of gene content variation with SNP distance for (A) 35 *S. flexneri* and (B) 23 *S. sonnei* pairs of isolates sampled at two-time intervals. Each dot represents a pair of isolates, by which the gene content variations between the isolates are plotted along the y-axis and the SNP distance between the isolates along the x-axis. Pairs are coloured by classification according to inlaid key which revealed difference in pattern among carriage and reinfection associated pairs, for both species. Spearman’s rank correlation coefficient value is displayed on the top right.

Then, we examined the effect of pair class (i.e. carriage or reinfection associated) on the level of accessory genome variation and disentangled the variations contributed by gain and loss events (Figure 2). Here, we define ‘gained’ as genes present in the later and absent in the earlier isolates of a pair, and vice versa for ‘lost’. This revealed the number of genes gained ranged from 5 to 93 (median = 21) and genes lost from 0 to 123 (median = 21) for *S. flexneri* carriage associated pairs (Figure 2). This was lower than the number of genes gained among *S. flexneri* reinfection associated pairs, which ranged from 9 to 213 (median = 82) and genes lost from 7 to 116 (median = 48). A similar relationship was seen for *S. sonnei*, where for carriage associated pairs, the number of genes gained ranged from 4 to 91 (median = 28) and genes lost from 4 to 176 (median = 24). For *S. sonnei* reinfection associated pairs, the number of genes gained ranged from 47 to 597 (median = 163) and genes lost ranged 57 to 182 (median = 141). For both species, the distribution of gene content variation between carriage associated pairs was significantly different to reinfection associated pairs for the number of genes gained (*S. flexneri p* = 0.11e-03 and *S. sonnei p* = 0.88e-03, Mann Whitney U test) and genes lost (*S. flexneri p* = 0.03 and *S. sonnei p* = 0.003) (Figure 2).

**Figure 2.**
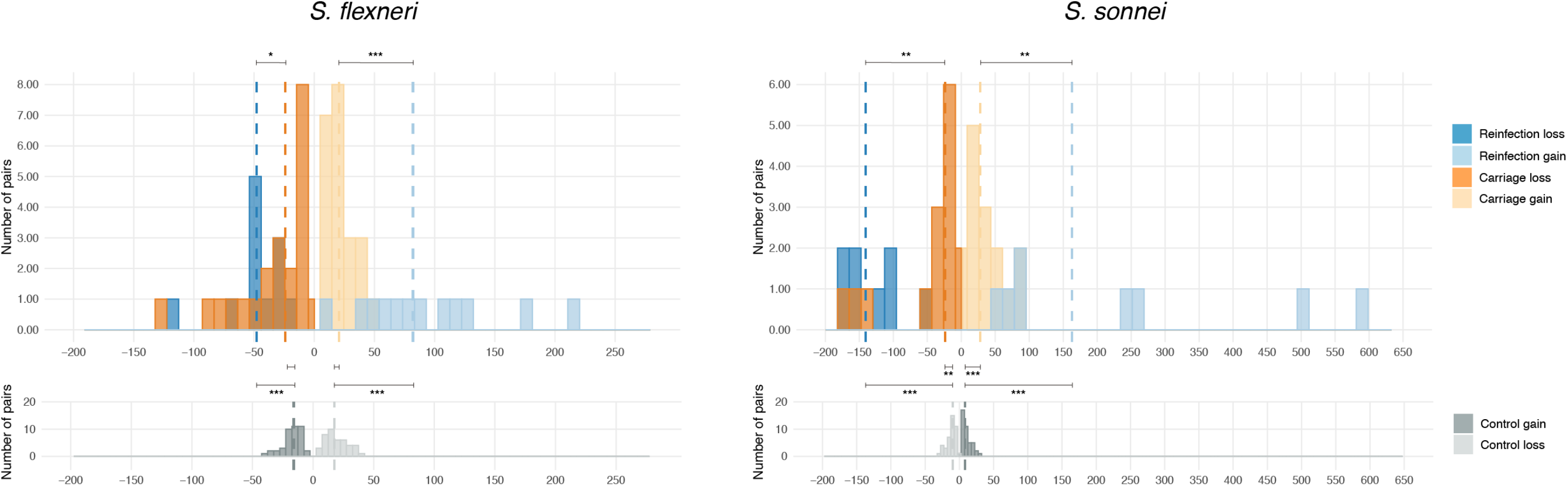
Distribution of the scale of accessory gene variation among paired isolates associated with carriage and reinfection in *S. flexneri* and *S. sonnei*. Frequency histogram plots show the number of accessory genes varying among isolate pairs. Genes present in the earlier serial isolate, but absent in the later are plotted as negative values (genes lost) and genes absent in the first, but present in the second isolate are plotted as positive values (genes gained). Variations derived from carriage or reinfection associated pairs are coloured according to the inlaid key, median values of distributions are shown as dashed vertical lines. Frequency histogram plots of *in silico* controls, showing intra-genome stochastic variation generated due to assembly, annotation and clustering are displayed below. Statistical differences between carriage and reinfection associated pairs, *in silico* controls and empirical data were tested using Mann-Whitney U tests, asterisks representing significance code * *p* < 0.05, ** *p* < 0.005 and *** *p* < 0.0005

As an important control for assessing whether the distribution of gene content variation for carriage associated pairs was biological in origin (rather than the result of stochastic variation in genome assembly, annotation and clustering) (Figure 2), we assembled genomes from synthetic read sets of varied length and insert size, generated from reference genomes, and performed pairwise homologous sequence comparison, similarly to above. Annotation of the synthetic genomes revealed variation in the number of coding sequences (CDS) ranged from 4215 – 4234 for *S. flexneri* 3a (20BP) and 4228 – 4247 for *S. sonnei* (Ss046) (Supplementary Table 3). Additionally, pairwise comparisons of the synthetic genomes generated substantial gene content variation (Figure 2). Specifically, for *S. flexneri,* the number of genes gained among *in silico* replicates of the same genome, ranged between 5 to 39 (median = 18) and genes lost between 5 to 39 (median = 16). For *S. sonnei* the number of genes gained ranged from 2 to 28 (median = 8) and genes lost from between 2 to 28 (median = 10). The gene content variation distributions generated from *in silico* genome replicates acted as controls and were statistically compared with the distribution among carriage pairs. This revealed a significant difference for the number of genes gained (*p* = 0.16e-03) and lost (*p* = 0.79e-03) between *S. sonnei* carriage associated pairs and the *in silico* control (Figure 2), indicating true biological variation between carriage associated pairs. Whereas for *S. flexneri*, there was no indication of statistically significant differences in genes gained (*p* = 0.74) or lost (*p* = 0.31) between carriage associated pairs and *in silico* controls, indicating that variations observed between isolates in carriage pairs were likely stochastic variations due to artefact. Gene content variation between reinfection associated pairs and *in silico* controls were significantly different (*p* ≤ 0.90e-05) for both *S. flexneri* and *S. sonnei* reinfection pairs (Figure 2).

### Gain/loss of AMR genes and known MGEs

As AMR is increasingly developing among *Shigella spp* in MSM, we screened for changes in genetic deteminants that confer resistance, including horitzontally aquired genes and point mutations. We also assessed for the presence of resistance determinants previously associated with MSM-associated *Shigella*. In particular, the presence of the pKSR100 plasmid which carries AMR genes conferring high-level resistance to azithromycin and associated with driving the success of MSM-associated *Shigella* sublineages (2, 4, 9). As expected, all *S. flexneri* and *S. sonnei* isolates within the dataset were multidrug resistant, habouring genetic determinants conferring resistance to three or more antimicrobial classes (Supplementary Table 4). When using short-read mapping to confirm presence of plasmids, we found the majority of *S. flexneri* isolates carried the *Shigella*-Resistance Locus Multi-Drug Resistance Element (SRL-MDRE) (64/68) and the pKSR100 plasmid (55/68), with only five isolates carrying the pCERC1 plasmid. For *S. sonnei*, all isolates carried the transposon Tn*7* and class II integrons (In2) with the majority (30/46) of isolates also carrying the spA plasmid and 43% (20/46) of isolates carrying the pKSR100 plasmid. The high AMR rates observed here reflect the known mobile genetic element content for UK *S. flexneri* and *S. sonnei*.

To look at changes in AMR over time, we explored what AMR genes were gained and lost over the course of *Shigella* infection. Here, we have applied the same working definition of gained and lost as previously mentioned. This revealed discrepancies in acquired AMR genes for 10 *S. flexneri* and 8 *S. sonnei* pairs, in line with population level trends (Table 2). Differences in AMR genes were observed between carriage and reinfection associated pairs for both species, often associated with the pKSR100 plasmid being acquired in reinfection pairs. Whereas, AMR genes associated with the pCERC1 plasmid were lost in carriage associated pairs. These individual trends of pKSR100 gain and pCERC1 loss are consistent with observations across MSM-associated shigellae (2, 9). Concerningly, there was evidence of AMR gain in two carriage associated pairs, with the extended beta-lactamase gene *blaSHV*_*12*_ being acquired by an *S. flexneri* 2a (Case ID I) pair and *blaTEM*_*1*_ in an S*. sonnei* pair (Case ID L) (Table 2), suggesting the possibility of AMR acquisition during persistent infection. A BLASTn search of the 52,219bp contiguous sequence carrying the *blaTEM*_*1*_ gene revealed 86% coverage and 99% identity with an *E. coli* O182:H21 plasmid (GeneBank accession: CP024250.1). The length of the contig carrying *blaSHV*_*12*_ spanned only the length of the gene, thus we were unable to reconstruct the full genetic context of this resistance gene and identify its origin.

**Table 2.** Variation in antimicrobial resistance genes detected among paired isolates of *S. flexneri* and *S. sonnei*.

| Reinfection/<br>Carriage | Species | Case<br>ID | Intervals<br>(days) | MDR<br>plasmid<br>gained* | MDR<br>plasmid<br>lost** | AMR genetic<br>determinants<br>associated |
| --- | --- | --- | --- | --- | --- | --- |
| Carriage | *S. flexneri*<br>3a | F | 27 |  | pCERC1 | *dfrA14*, *sul2*,<br>*strA/B* |
|  |  | O | 83 |  | pCERC1 | *sul2*, *strA/B* |
|  | *S. flexneri*<br>2a | I | 6 |  |  | *blaSHV-12* |
|  | *S. sonnei* | L | 35 |  |  | *blaTEM-1* |
| Reinfection | *S. flexneri*<br>2a | A | 1142 | pKSR100 |  | *mph(A)*, *blaTEM-1*,<br>*dfrA17*, *sul1*,<br>*aadA5* |
|  |  | C | 496 |  | pCERC1 | *dfrA14*, *sul2*,<br>*strA/B* |
|  | *S. flexneri*<br>3a | C | 805 | pKSR100 |  | *erm(B)*, *mph(A)*,<br>*blaTEM-1* |
|  |  | E | 193 |  | pKSR100<br>integron | *dfrA17*, *sul1*,<br>*aadA5* |
|  |  | I | 1862 | pKSR100 |  | *erm(B)*, *mph(A)*,<br>*blaTEM-1* |
|  |  | D | 1099 |  | pCERC1 | *dfrA14*, *sul2*,<br>*strA/B* |
|  |  | J | 905 | pKSR100 | pCERC1 | *mph(A)*, *blaTEM-1*,<br>*dfrA17*, *sul1*,<br>*aadA5*, *dfrA14*,<br>*sul2*, *strA/B* |
|  | *S. sonnei* | B | 925 | spA |  | *strA/B*, *sul2*, *tetA* |
|  |  | C | 1409 | spA,<br>pKSR100 |  | *strA/B*, *sul2*, *tetA*,<br>*blaTEM-1*, *erm(B)*,<br>*aadA5*, *dfrA17*,<br>*sul1*, *mph(A)* |
|  |  | D | 42 |  | spA | *strA/B*, *sul2*, *tetA* |
|  |  | G | 1208 | spA,<br>pKSR100 |  | *strA/B*, *sul2*, *tetA*,<br>*blaTEM-1*, *erm(B)*,<br>*aadA5*, *dfrA17*,<br>*sul1*, *mph(A)* |
|  |  | I | 659 | pKSR100 |  | *blaTEM-1*, *erm(B)*,<br>*mph(A)* |
|  |  | J | 481 | pKSR100 |  | *blaTEM-1*, *erm(B)*,<br>*mph(A)* |
|  |  | V | 184 |  |  | *aadA1* |

For reinfection associated pairs, \*plasmid gained is defined as the plasmid being present in the second isolate but absent in the first isolate of the pair, and \*\*plasmid lost is defined as the plasmid being absent in the second isolate but present in the first isolate of the pair.

Point mutations in the QRDR were identified in 29/46 (63%) *S. sonnei* isolates, 19 of which were triple mutations (*gyrA* S83L, *gyrA* S87G, *parC* S80I) known to confer resistance against ciprofloxacin, and 10 with single mutation (*gyrA* S83L and D87G) conferring reduced susceptibility (Supplementary Table 4). Single *gyrA* S83L mutations were detected in two *S. flexneri* 3a isolates. Although the rates of quinolone resistance were moderate in *S. sonnei* and low in *S. flexneri*, there was no sign of *de novo* mutation in the QRDR region over the course of infection as we did not observe any isolate pairs with the same genotype (i.e carriage associated pairs) acquiring mutations in later isolates.

### Generation of an important MSM-associated S. flexneri 3a isolate reference genome and the establishment of carriage/reinfection associated pairs

To determine structural variation and genome rearrangement of *S. flexneri* over time, we PacBio sequenced 16 isolates from an epidemic sublineage of MSM-associated *S. flexneri* 3a, serially isolated from eight individuals at time intervals of 9 to 911 days apart (Figure 3A and 3B). Notably, Illumina data from these isolates were already available from a previous study (2). In the process, a complete genome for isolate 20BP was generated, which comprised of a chromosome of 4,522,047bp, the virulence plasmid of 231,165bp and the pKSR1000 plasmid of 72,593bp. This complete genome was used as a high-quality reference genome for further analyses and has been deposited under accession number GCA_904066025 in NCBI.

**Figure 3.**
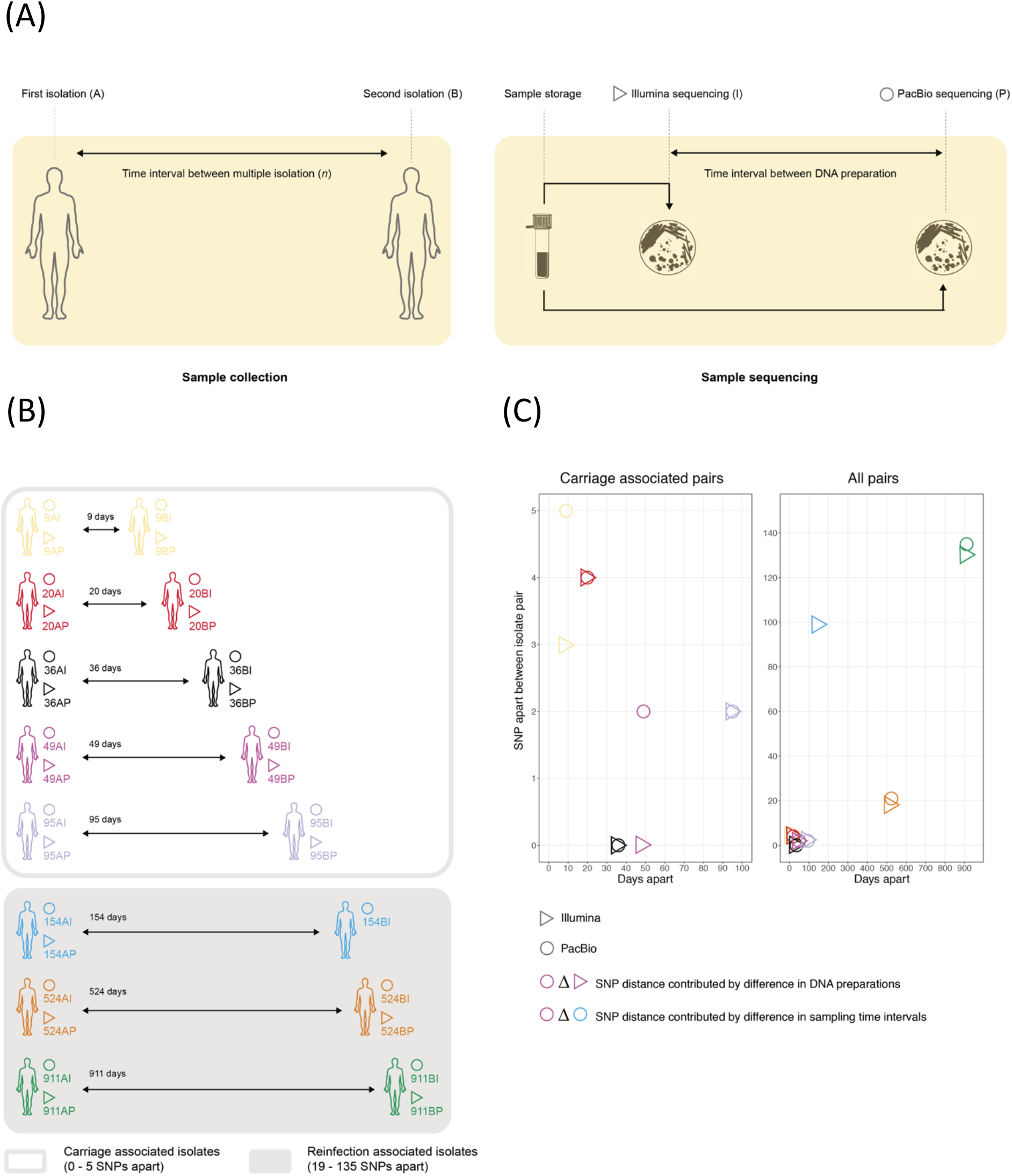
Sampling time points and sequencing technologies used to investigate large structural variation of *S. flexneri* 3a genomes over time. (A) Isolates from each pair were serially sampled from the same patient. Following data collection, the samples were stored and later Ilumina sequenced. After a considerable amount of time (e.g. several years), the samples were revived and PacBio sequenced. (B) Serial isolation of *S. flexneri* 3a was performed across 8 patients sampled between 9 and 911 days apart. Names of genome data used in the current study are presented in the diagram according to its abbreviation. (C) The plots display SNP distance and days apart between serial sampling of the *S. flexneri* 3a isolate pairs. Each shape on the plot represents variations between an isolate pair. The colour of each shape represents the time interval between sampling of an isolate pair, as demonstrated in (B). Different shapes represent the sequencing technology used according to the inlaid key. Plot on the left contain carriage associated pairs and plot on the right contain all isolate pairs analysed in this study. This revealed distinctive clustering, by which pairs associated with carriage are isolated at shorter time interval with less SNPs apart than compared to reinfection associated pairs.

Genetic distances of the eight isolate pairs sampled at various time intervals ranged from 0 to 135 SNPs apart (Figure 3C). Generally, SNP distances between pairs increased with time interval between serial isolations. This was consistent with the previously established epidemiological definitions, and the same definition of carriage and reinfection associated pairs was applied (see methods). As PacBio sequencing of isolate 154BP failed (Figure 3B), there were in total long-read sequenced genomes for seven serial isolate pairs; five carriage and two reinfection associated pairs. SNP distances between replicate sequencing of individual isolates using PacBio and Illumina (e.g. between 9AI and 9AP) were equivocal, with variation contributed by different sequencing preparations being 0 to 5 SNPs apart (Figure 3C).

### Large-scale variation of S. flexneri genome over time

To detect structural rearrangements among the seven pairs of *S. flexneri* 3a, we aligned all PacBio sequenced genomes against the high-quality reference genome of 20BP and assessed discrepancies between each pair. We identified a total of 34 structural variations in the 7 pairs of isolates across 14 genomic regions, including 9 copy deletions, 7 insertions, 7 duplications, 5 inversions, 4 deletions, 1 translocation and 1 translocation inversion (Figure 4A). Three structural variants were less than 1,500bp and mapped to IS elements. We analysed sequences at the borders of the remaining 31 variants to determine possible mechanisms facilitating the rearrangements. This revealed 15 variants had occurred through recombination between homologous IS copies and two variants had occurred through recombination between ribosomal operons (Supplementary Table 5). Of the remaining 14 variants, 7 possessed IS sequence in only one end. We did not detect presence of repeat sequences or IS elements at the borders of the remaining 7 variants, thus rearrangements have been facilitated by an unknown mechanism.

**Figure 4.**
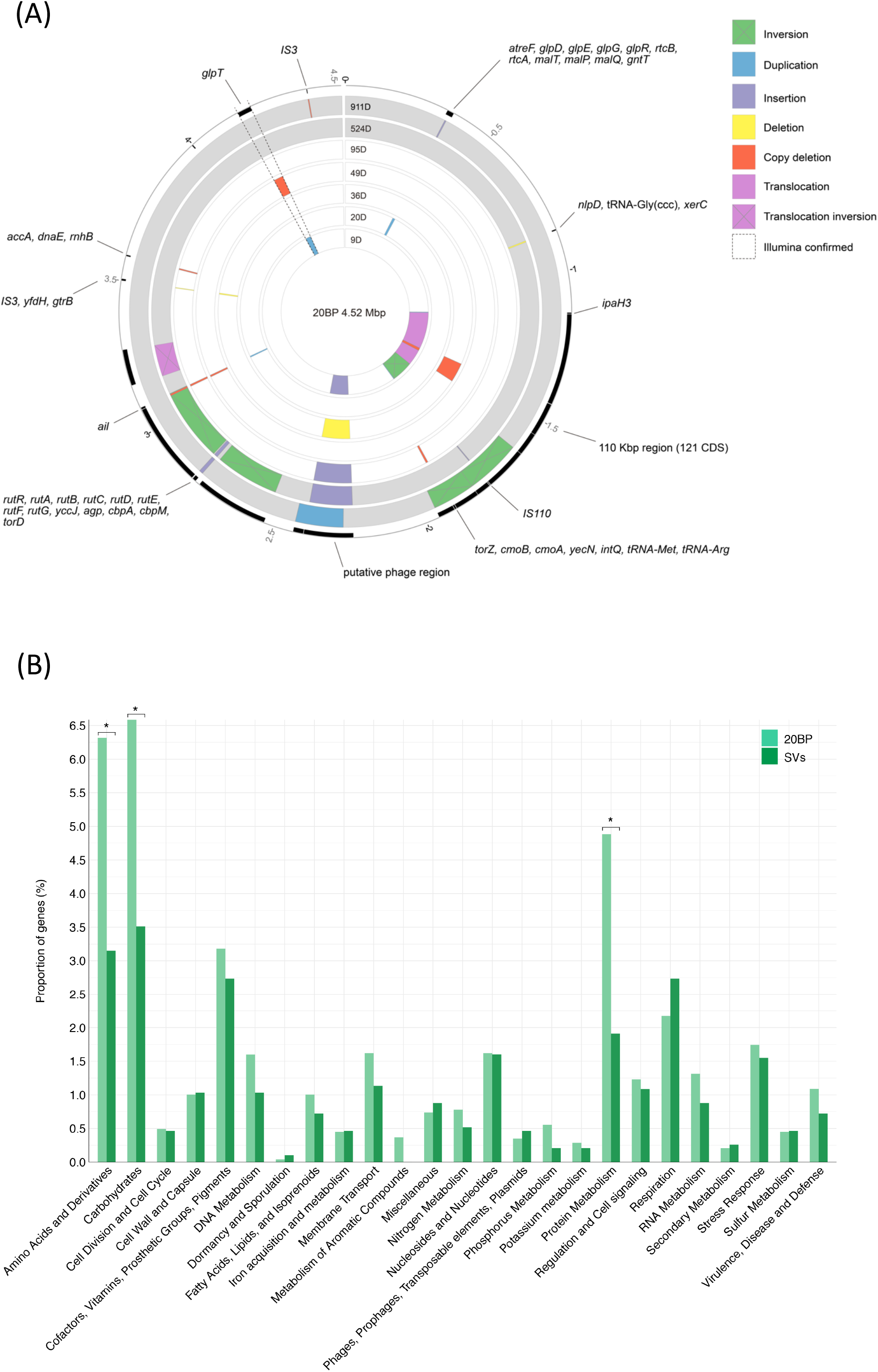
Chromosomal structural variation detected among seven pairs of serially-isolated MSM-associated *S. flexneri* 3a isolates and functional annotation for genes and prediction of the GO category by RAST. (A) The concentric rings represent pairwise comparisons between PacBio generated genomes, with the time interval in days overlaid at uppermost and increasing in out circles. Genomes of two isolate pairs associated with reinfection (rather than carriage) are highlighted in grey. Overlaid are coloured blocks according to the inlaid key indicating the nature and frame of structural variations. The outermost track displays predicted genes within regions of structural variations (genes encoding for hypothetical proteins are not annotated). Duplication and deletion events confirmed by both Illumina and PacBio data are highlighted in dashed lines. (B) The proportion of genes in GO categories annotated by RAST for 20BP reference chromosome (light green bars) and structural variable regions (dark green bars). GO categories which have significant (*p* < 0.05) difference in proportion of genes are indicated by an asterisk. Genes unable to be assigned to a category are not displayed, which represented 59% for 20BP reference chromosome and 72% for regions of structural variation.

A total of 1,791 genes were found within the genomic regions of structural variations and rearrangements. In order to see if particular gene functions were enriched within these genomically plastic regions, we annotated all genes across the 20BP chromosome and assigned them to predicted functional categories according to GO categories (see methods). A chi-square test was used to compare the number of genes across the variable regions and the total number of genes across the entire reference chromosome for each category. Overall, there was significant difference in the number of genes belonging to 3 categories (Figure 4B). Specifically, genes predicted with function in the amino acids and derivatives, carbohydrates and protein metabolism GO categories were significantly depleted in variant regions.

Rearrangements occurring in two regions were commonly observed, including a 166 kbp region at ~2.24 – 2.40 Mbp and an 8 kbp region at ~3.07 – 3.08 Mbp identified among four pairs (3 carriage and 1 reinfection) (Figure 4A). The former region is flanked by *IS91* copies, carries 207 predicted genes and encodes an incomplete prophage. The reinfection associated pair sampled 911 days apart displayed duplication of an overlapping region offset by 37 kbp to the incomplete prophage, also flanked by *IS*91 copies (Figure 4A). Regarding the second region, the 8 kbp region falls within an intact prophage, flanked by homologous *IS1* copies and contains 11 genes. The majority (10/11) of which encodes for hypothetical proteins and a predicted Ail/Lom family outer membrane β-barrel protein. By which, the bacterial Ail protein is a known virulence factor thought to promote host cell invasion (39), and Lom is a phage protein expressed during lysogeny (40).

In order to confirm deletion and duplication events as biological and not as an effect of different sequencing preparations, we performed short read mapping of the Illumina sequenced data against the 20BP reference genome, which confirmed variation for only one duplicated/deleted region of 127kbp at 4.17 – 4.22 Mbp (dashed lines, Figure 4A) flanked by rRNA operons at the borders. This region varied in two carriage pairs, with duplication of this region being observed in the pair sampled 9 days apart and deletion of the same region observed in the pair sampled 49 days apart (Supplementary Figure 3). The region contains 37 CDS, including *ompA* which encodes the outer membrane protein A, a virulence factor involved in facilitating cell-to-cell spread and a target for vaccine development (41, 42). Although consistency between Illumina and PacBio data indicates a region that genuinely changed over the course of infection, this was not in a uniform direction over time and the lack of confirmation of other regions suggests that some structural variations detected may have arisen either from different preparations or occurred during sample storage, which was of considerable duration.

## Discussion

In the current study, we characterised and compared the accessory genome dynamics of *S. flexneri* and *S. sonnei* isolates associated with both carriage and reinfection, as defined previously based on SNP typing data and time intervals between serial-isolation (17). Our results reveal that carriage and reinfection pairs differ, and that SNP distance and the magnitude of gene content variation correlated, albeit the association was weaker for *S. flexneri*. In general, reinfection associated pairs had greater SNP-distance and varied by a greater number of genes than carriage pairs.

The dynamics of accessory genes between carriage and reinfection associated pairs were examined and showed that for both species, there was a significant difference in gene content variation between the two pair classes. This supports the concept of a decreased genetic distance in carriage compared with reinfection associated pairs. Although we have used working definitions of carriage and reinfection associated within the manuscript to narrate our study, it is important to note that further clinical and epidemiological data (which is unavailable) would be required to fully differentiate between persistent carriage or chronic infection, and reinfection with closely and more distantly related isolates.

*Shigella* accessory genomes are highly plastic, and the acquisition of AMR genes through horizontal gene transfer (HGT) have been previously shown to enhance and drive MSM-associated shigellae epidemics (9, 10). Here, we assessed the gain and loss of AMR genetic determinants conferring resistance in carriage and reinfection pairs to investigate acquisition of resistance during persistent infection. We detected acquisition of different extended beta-lactamase genes in *S. flexneri* and *S. sonnei* carriage associated pairs. Although we were unable to conclude the origin of both genes, acquisition of AMR genes in *Shigella* have been previously speculated to be facilitated through transfer of plasmid between *Escherichia coli* within the gut (43–45) and an identical multidrug resistance plasmid isolated from *S. sonnei* and *E. coli* in a single patient has been reported (46). Thus, acquisition of the AMR genes within the two carriage pairs within this dataset indicates possible HGT occurring during persistent infection, possibly from *E. coli*. And, while acquisition of AMR through HGT in hospital settings has been documented (47), here we have observed AMR gain in patient infections in a community setting.

Structural variations and genome rearrangements have played an important role in the evolution of *Shigella* (48). Thus, aside from changes in the accessory genome it is also important to consider what larger structural variations and rearrangements may occur during *Shigella* infection over time. To do this, we examined the genomes of 16 (seven pairs) MSM-associated *S. flenxneri* 3a epidemic sublineage isolates serially sampled at various time intervals and identified numerous variations and rearrangements. Few regions of the chromosome demonstrated different types of variation at the same location among different pairs. These all had IS elements or rRNA operon at the borders, which most likely facilitated the variation at these regions (49, 50). Functional prediction of genes located within the structural variant regions revealed depletion of genes involved in key metabolic processes including amino acids and derivatives, carbohydrates and protein metabolism. Since large rearrangements can be deleterious (51), it is evident that these genes may be functionally important for *Shigella*, as they were less prone to structural variation and rearrangements (52). Mapping of Illumina data confirmed genuine duplication and copy deletion in two carriage associated pairs at a 127 kbp region, which carries the *ompA* virulence factor. Although parallel variation demonstrates genomic instability of this region, we do not know if this has an impact on the virulence phenotype.

As well as detecting genuine biological variation that occurred over the course of infection, we detected many structural variations that were artefactual. This may have been from the impact of distinct sample preparations, or, more likely given that the variations occurred in common regions across isolates, may have arisen from adaptation/changes during prolonged storage. The most prominent category of artefactual variation was the deletion of a 166 kbp prophage region in six PacBio sequenced genomes. Illumina data generated from different DNA preparation revealed this region was in fact present in the original isolates. Since there was considerable duration between the DNA preparations, the loss of this region exclusively in the PacBio sequenced genomes could be due to the loss of selection for genes whose function are no longer required within the storage environment, whereby genes with dispensable function are discarded (53, 54). Such events have been shown to play an important part in the convergent evolution of *Shigella* species as a host-restricted pathogen (48). The deletion of this region under the storage environment but retention in the clinical environment suggests genes in this region may have functions contributing to infection and/or ecological interaction, and thus warrants further investigation. Furthermore, this highlights the importance of well stored samples for the inclusion in studies and due caution when examining large-scale genomic rearrangements.

In summary, we have utilised isolate pairs occurring in a comparatively new infection setting for *Shigella* to characterise the accessory genome dynamics that occur in persistent infection (and contrasted this with reinfections). We showed an overall gain of AMR across isolate pairs, consistent with population trends of AMR among MSM-associated shigellae. We also detected structural variation in carriage associated pairs over time and found that some structural changes were the result of storage/preparation artefact, both of which may have biological relevance. It is worth noting that due to the limited sampling intervals and methodology applied, we will not have captured all possible variations (i.e transient, small and single gene variant). However, we have provided novel insights to large structural chromosomal variations in *Shigella* over time, and this is an important step in trying to understand how the pathogen might adapt during persistent infection. Finally, our study additionally highlights the need for appropriate controls in genome studies and the storage of high-quality reference isolates. To that end, we have deposited the cognate strain for the 20PB reference genome of the intercontinentally transmitting *S. flexneri 3a* in NCTC.

## Acknowledgements

This work was supported by a UKRI MRC NIRG award (MR/R020787/1) and a technology directorate voucher from the University of Liverpool. KSB is supported by a Wellcome Trust Clinical Research Career Development Award (106690/A/14/Z). The authors are grateful to Sam Haldenby, Matthew Gemmell and Richard Gregory at the Centre for Genomics Research, University of Liverpool for technical support.

---

**Table S3.**
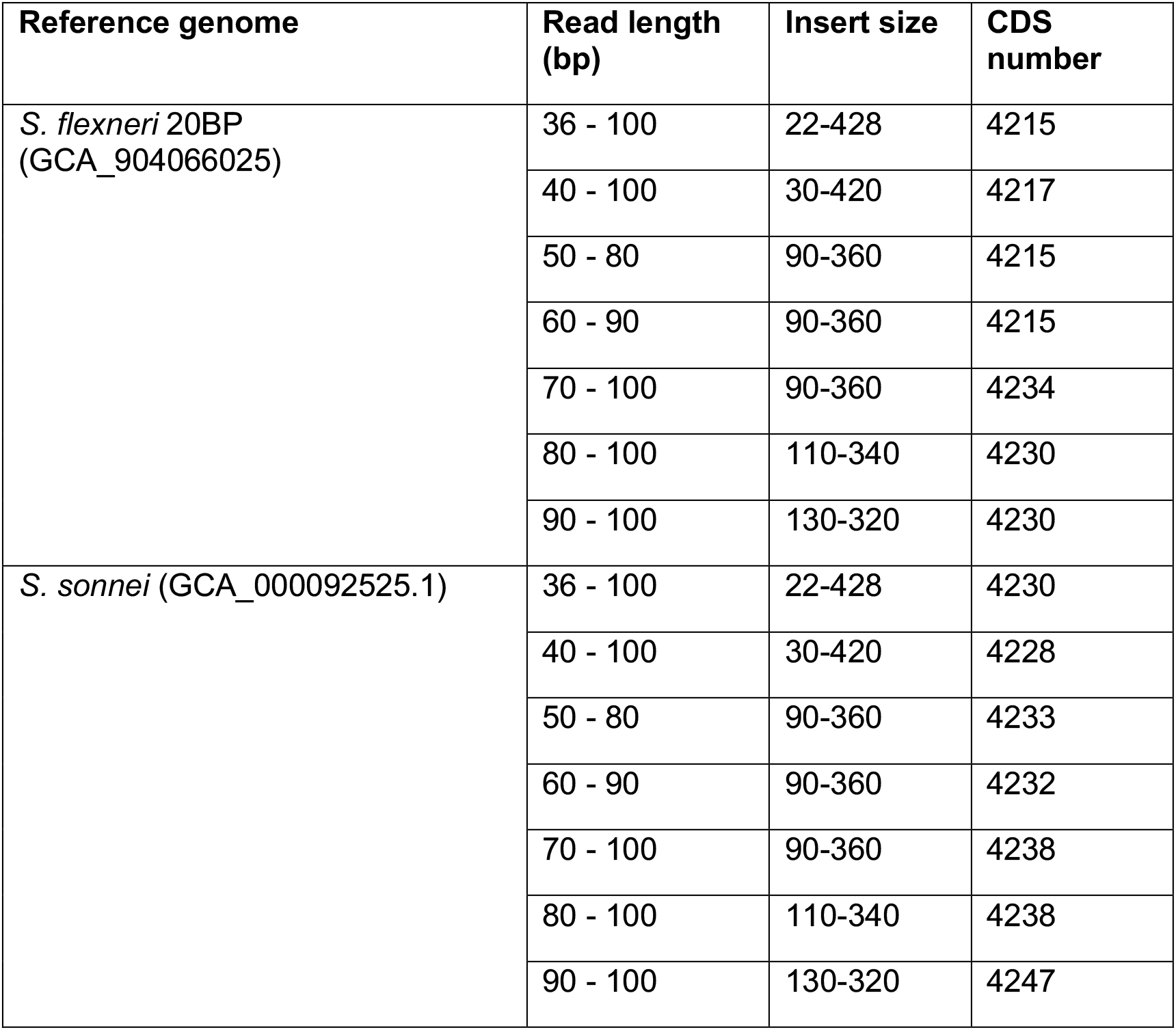
Number of CDS annotated from draft genome assemblies generated from synthetic reads of various length and insert size.

| Reference genome | Read length (bp) | Insert size | CDS number |
| --- | --- | --- | --- |
| *S. flexneri* 20BP (GCA_904066025) | 36 - 100 | 22-428 | 4215 |
|  | 40 - 100 | 30-420 | 4217 |
|  | 50 - 80 | 90-360 | 4215 |
|  | 60 - 90 | 90-360 | 4215 |
|  | 70 - 100 | 90-360 | 4234 |
|  | 80 - 100 | 110-340 | 4230 |
|  | 90 - 100 | 130-320 | 4230 |
| *S. sonnei* (GCA_000092525.1) | 36 - 100 | 22-428 | 4230 |
|  | 40 - 100 | 30-420 | 4228 |
|  | 50 - 80 | 90-360 | 4233 |
|  | 60 - 90 | 90-360 | 4232 |
|  | 70 - 100 | 90-360 | 4238 |
|  | 80 - 100 | 110-340 | 4238 |
|  | 90 - 100 | 130-320 | 4247 |

**Supplementary Figure 3.**
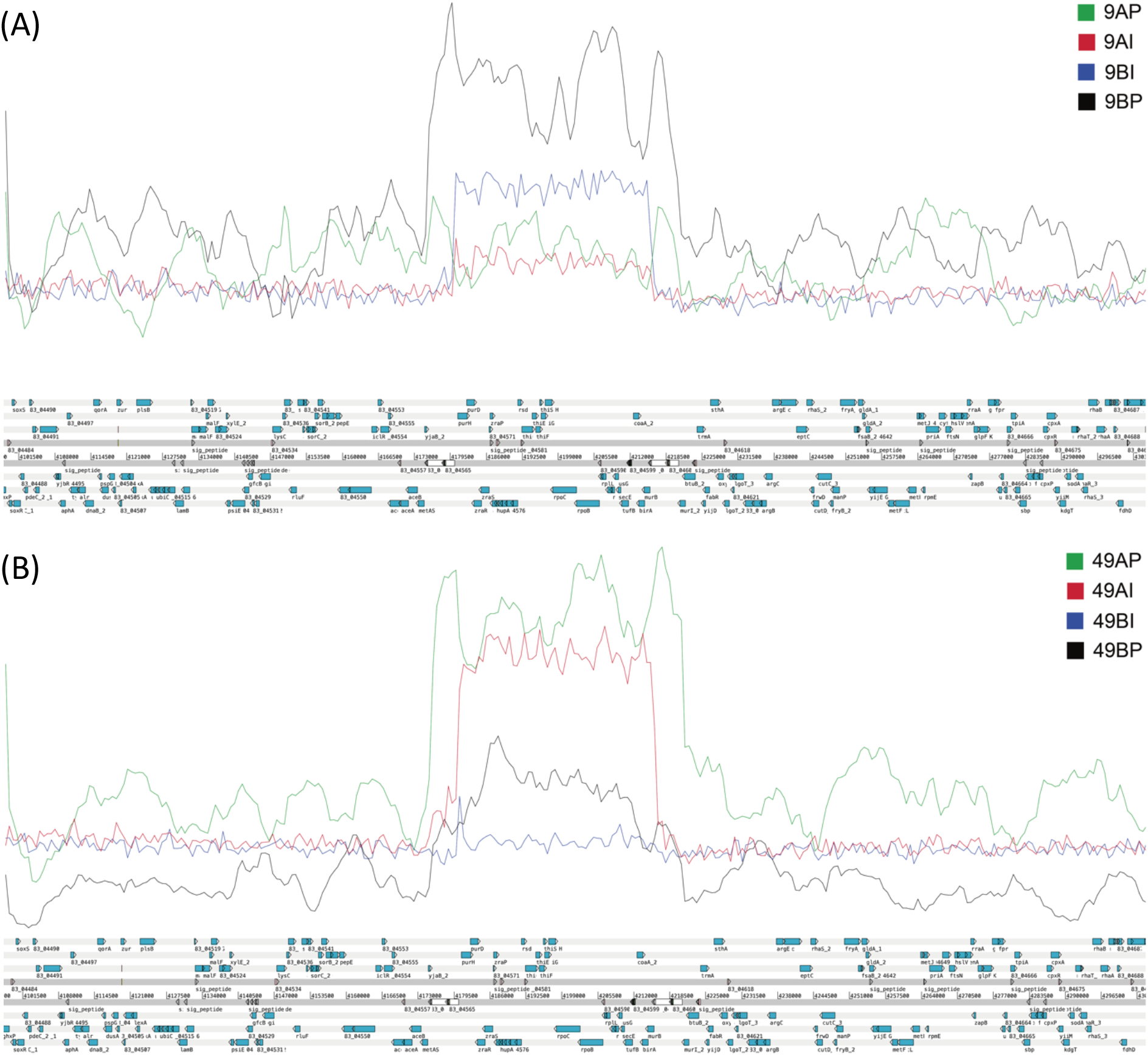
Mapping of short and long reads of a 127 kbp region at 4.17 – 4.22 Mbp confirming duplication/deletion events. (A) Reads of isolates sampled at 9 days interval were mapped against complete reference sequence of 20BP, which confirmed duplication of the region in the isolate sampled at the second time interval. (B) Reads of isolates sampled at 49 days interval, confirmed copy deletion of the same region in the isolate sampled at the second time interval.

